# Exercise-induced plasma-derived extracellular vesicles increase adult hippocampal neurogenesis

**DOI:** 10.1101/2025.08.02.668291

**Authors:** Meghan G. Connolly, Alexander M. Fliflet, Prithika Ravi, Dan I. Rosu, Marni D. Boppart, Justin S. Rhodes

## Abstract

Aerobic exercise enhances cognition in part by increasing adult hippocampal neurogenesis, angiogenesis, and astrogliogenesis. Since hippocampal atrophy is a hallmark of several neurological and psychiatric conditions—including depression, PTSD, Alzheimer’s disease, and aging—understanding the mechanisms by which exercise increases neurogenesis has broad therapeutic relevance. One potential mechanism involves extracellular vesicles (EVs), lipid bilayer-enclosed particles released by multiple tissues during exercise that transport bioactive molecular cargo to distant organs, including the brain. In this study, we tested whether plasma-derived EVs from exercising mice (ExerVs) are sufficient to promote hippocampal neurogenesis, astrogliogenesis, and vascular density in sedentary mice. EVs were isolated from the plasma of sedentary or exercising C57BL/6J mice and injected intraperitoneally into sedentary recipients twice weekly for four weeks. To evaluate reproducibility, the study was conducted across two independent cohorts using identical procedures. ExerV-treated mice showed a significant increase in BrdU-positive cells in the granule cell layer compared to both PBS- and SedV-treated controls in both cohorts. Approximately 90% of these cells co-expressed NeuN, indicating neuronal differentiation, while 6% co-expressed S100β, indicating astrocyte generation. No changes were observed in vascular density across groups. These findings provide initial evidence that systemically delivered exercise-derived EVs can enhance hippocampal neurogenesis and astrogliogenesis in sedentary mice. This proof-of-concept work supports further investigation into ExerVs as a potential therapeutic strategy for conditions associated with hippocampal atrophy.

**Highlights:**

- ExerVs increase hippocampal neurogenesis and astrogliogenesis in sedentary mice
- ExerVs do not affect hippocampal vascular area
- Circulating ExerVs are sufficient to recapitulate key brain benefits of exercise

## 1. Introduction

Aerobic physical exercise is a highly effective method for preserving cognitive function across the lifespan^1–3^. Understanding the mechanisms by which exercise enhances cognitive abilities holds profound implications for overall well-being, especially considering the prevalent issue of cognitive decline during the natural aging process^4^. Numerous studies have documented enduring neurological alterations resulting from aerobic exercise training, which are strongly correlated with observed enhancements in cognitive performance^5–7^. In particular, long-term modifications within the hippocampus have been consistently reported in response to aerobic exercise training, indicative of bolstered cognitive function^1,6^. For instance, in rodent models, voluntary wheel running increases adult neurogenesis in the dentate gyrus of the hippocampus by two-to five-fold, depending on genotype and exercise intensity^8,9^. Understanding how exercise increases neurogenesis could have important therapeutic implications. Nevertheless, despite decades of study, the mechanisms responsible for exercise-induced neurogenesis remain largely unknown.

Research over the past few decades has suggested that bloodborne factors released from skeletal muscle and other organs cross into the brain leading to increased proliferation and survival of new neurons in the hippocampus^10–14^. Indeed, studies have shown that the beneficial effects of exercise can be transferred through injections of plasma from voluntarily running animals into aged mice causing increased hippocampal neurogenesis and improved cognitive function^15^, likely through decreased inflammation^16^. Multiple bloodborne factors have been implicated including vascular endothelial growth factor (VEGF)^17^, insulin-like growth factor-1 (IGF-1)^18^, platelet factor 4 (PF4)^14^, selenoprotein P (SELENOP)^19,20^, peroxisome proliferator-activated receptor gamma coactivator-1 alpha (PGC-1α)/FNDC5/irisin^13^, cathepsin B^11^, L-lactate^21^, and interleukin (IL)-6^22,23^. While these molecules have been shown to be involved in exercise-induced neurogenesis, they likely represent only part of a broader, more coordinated signaling system through which exercise exerts its effects on the brain.

Additional factors such as circulating extracellular vesicles (EVs) that contain multiple different molecules could be important signaling vehicles from the periphery to the brain. EVs carry different molecular cargo depending on their cellular biogenesis, but they are generally comprised of various proteins, lipids, DNA, RNA, and microRNAs (miRNAs)^24–26^. Literature has shown that during exercise, EVs are released at a greater rate from various sources, including skeletal muscle into circulation^27–29^. Indeed, Frühbeis et al. (2015) showed that EVs were released into the circulation during an early phase of exercise, before the individual anaerobic threshold. Guescini et al. (2015) demonstrated that these EVs released into circulation may originate from skeletal muscle by demonstrating that the systemic EVs produced during exercise were positive for the muscle-enriched α-sarcoglycan protein and contained high levels of the skeletal muscle-specific microRNA, miR-206.

EVs are present in various biofluids and both *in vitro* and *in vivo* studies have demonstrated that EVs can traverse the blood-brain barrier, making them promising candidates for enacting changes in the central nervous system^30–33^. Furthermore, there is growing evidence indicating that EVs, especially their contents, contribute to various brain functions such as synaptic plasticity, intercellular communication, and neurogenesis^34–36^. For example, intranasal administration of EVs derived from neural stem cells (NSCs) derived from human induced pluripotent stem cells into adult rats and mice promoted hippocampal neurogenesis^37^. Together, the literature indicates that EVs are important molecules released during exercise and they can exert changes in the hippocampus. However, whether EVs taken from circulation following exercise are sufficient to increase adult hippocampal neurogenesis remains unknown.

The present study sought to determine whether plasma-derived extracellular vesicles (EVs) from exercising mice (ExerVs) are sufficient to enhance hippocampal neurogenesis, and whether these effects are accompanied by changes in astrogliogenesis and hippocampal microvasculature, key components of the neurogenic niche^38–42^. Multiple studies have shown that voluntary running enhances astrogliogenesis and vascular density in conjunction with neurogenesis ^38–40,43,44^. Astrocytes provide structural and metabolic support to newborn neurons and are highly responsive to circulating factors, particularly those released by contracting muscles^39,40,43–45^. Our lab previously showed that electrically stimulated muscle contractions increased astrocyte proliferation in the hippocampus, suggesting a potential muscle-to-astrocyte signaling axis^45^. This effect was recapitulated in an *in vitro* study applying isolated muscle cell factors to primary hippocampal neuronal cultures^46^.

A companion paper, studying the peripheral tissues of the same mice used herein reported a positive effect of ExerVs on vascular density in the gastrocnemius muscles^47^. That paper also reported the detailed methods and validation of the EVs. Further the paper reported on proteomic profiling of the ExerVs relative to EVs collected from sedentary mice (SedVs). Results revealed ExerVs displayed enrichment of proteins associated with neuroplasticity, cellular signaling, and synaptic remodeling. For example, ExerVs displayed increased levels of Spp1, a secreted glycoprotein known to influence cell survival, neuroinflammation, and adult hippocampal neurogenesis^48,49^. ExerVs also displayed increased Msn, a cytoskeletal regulator implicated in neural development and synapse formation^50,51^, and increased Psma3 and Psmb2, core proteasome subunits involved in protein degradation and turnover of synaptic proteins implicated in exercise-induced hippocampal neurogenesis^52,53^. Based on these data, we hypothesized that ExerVs, when administered to sedentary mice, would increase adult hippocampal neurogenesis, astrogliogenesis and microvascular area fraction.

## 2. Materials and Methods

### 2.1 Experimental Subjects

A total of 75 adult male C57BL/6J mice (4-month-old) were purchased from Charles River (Wilmington, MA, USA) and Jackson Laboratory (Jackson, ME, USA) and maintained in ventilated cages with *ad libitum* access to food and water throughout the experiments. Donor mice (described below) were singly housed while recipient mice (described below) were group housed. The colony room was maintained at on a 12-hour light/dark cycle. The experiments were approved by the University of Illinois Urbana-Champaign Institutional Animal Care and Use Committee (IACUC) (#22062) and followed the Guide for the Care and Use of Laboratory Animals.

### 2.2 Experimental Design

#### EV Donors

Mice used as extracellular vesicle (EV) donors (n=40 across two cohorts) were housed individually with access to horizontal running wheels (Low-Profile Wireless Running Wheel, Med Associates Inc., Fairfax, VT, USA) for 28 days. Wheels were either unlocked to allow voluntary running (exercise group, n=20) or locked to prevent movement (sedentary controls, n=20). Running distance was automatically recorded. Mice were fasted overnight and euthanized immediately following their final bout of exercise (approximately 3:00 AM). Blood was collected via cardiac puncture into 2 mL low-protein binding tubes and centrifuged at 4000 × g for 15 minutes to obtain platelet-free plasma. EVs were subsequently isolated from pooled plasma as described below.

#### EV Recipients

Recipient mice (n=29) were divided into two independent cohorts to confirm the reproducibility of observed effects. Findings from the first cohort prompted replication in Cohort 2 using identical treatment conditions. Based on results from Wu et al. (2023), 3×10^8^ particles of EVs were intraperitoneally (i.p.) injected twice per week for 4 weeks. Intraperitoneal (i.p.) injections were used to deliver EVs, as this route has been shown to result in reliable systemic uptake and detectable biodistribution across multiple organs, including the brain, while avoiding the technical challenges associated with intravenous injection ^55^. To label dividing cells, mice also received intraperitoneal injections of 5-bromo-2’-deoxyuridine (BrdU; 50 mg/kg) once daily for the first 10 days of the experiment, excluding weekends. The following treatments were injected: PBS as a vehicle control (n=10), sedentary EVs (SedV) as a condition control (n=9), and exercise EVs (ExerV) (n=10).

Mice were euthanized 24 hours after the last injection, and final body weights were recorded. For euthanasia, mice were anesthetized with an i.p. injection of sodium pentobarbital and brains were collected following transcardial perfusion with 0.9% saline. Brains were collected and postfixed overnight in 4% paraformaldehyde (4°C) then stored in 30% sucrose (4°C) until sectioning. A cryostat was used to section brains into 40 μm coronal sections and sections were stored (−20°C) in cryoprotectant solution.

### 2.3 Extracellular Vesicle Isolation and Characterization

The methods for EV isolation, characterization, and injection were aligned with those described in Wu et al. (2023). EVs were isolated via size exclusion chromatography (SEC) (Fractions #4-6 or Fractions #7-10; qEV single or original/35 nm, Izon Science, Medford, MA, USA) and ultrafiltration (Amicon® Ultra-2 Centrifugal Filter Unit – 30 kDa cutoff; Millipore, St. Louis, MO, USA). Additionally, the protein fraction (#8-10 qEV single or #16-25 qEV original) from SEC columns was also collected for use as an EV-depleted control. Nanoparticle tracking analysis (NTA; NanoSight NS300, Malvern Panalytical, Malvern, United Kingdom) was used to evaluate EV concentration and size, and EVs were verified via nanoLC-MS and CryoEM. Recent research has indicated that EVs tend to aggregate in the freezer and their concentration decreases with long-term storage^57,58^. Consequently, EVs were aliquoted (3×10^8^ particles/injection) and temporarily stored at −80°C until injection.

### 2.4 Immunohistochemistry and Image Analysis: Cell Genesis and Microvascular Density

To evaluate cell genesis, a one-in-six series from each of the EV recipient groups were stained for BrdU. Free-floating sections were first washed in tris-buffered saline (TBS), followed by 30 minutes in 0.6% hydrogen peroxide, and then again washed in TBS. For antigen retrieval, sections were first incubated in 50% deionized formamide (65°C for 90 minutes), followed by 2X saline-sodium citrate buffer (room temperature for 15 minutes), then 2N hydrochloric acid (37°C for 30 minutes), and finally in borate buffer (room temperature for 10 minutes). Sections were then blocked with 5% goat serum and 0.1% Triton-X in TBS for 30 minutes and then incubated in the rat anti-BrdU (1:200; AbCam Cat# ab6326) primary antibody for 48 hours at 4°C. After 48 hours, sections were washed, exposed to a second blocking step, and incubated in a biotinylated goat anti-rat (1:250; Vector Laboratories Cat# BA-9401-.5) secondary antibody (room temperature for 1 hour and 40 minutes). Sections were then incubated in avidin-biotin complex buffer (Vector Laboratories, Burlingame, CA) for one hour and then stained with the chromogen diaminobenzidine (DAB) (EMD Millipore Cat# D4418-50SET). To evaluate microvasculature, the staining procedure was similar to the cell proliferation procedure except antigen retrieval was achieved using a 25% pepsin (AbCam Cat# ab64201) solution (37°C for 10 minutes). Microvasculature was stained using a rabbit anti-Collagen IV (1:300; Bio-Rad Laboratories Cat #2150-1470) primary antibody and a goat anti-rabbit (1:250; Vector Laboratories Cat# BA-1000-1.5) secondary antibody.

Images of the granular cell and molecular layer were collected from 8-9 sections per animal on a Zeiss light microscope at 10X magnification. BrdU-positive cells were captured from a single plane selected through the z-axis. Estimates of the number of positive cells were hand counted and the volume of the granular cell layer (GCL) was obtained using Fiji software. The granular cell layer and molecular layer were outlined in Fiji and the area in pixels was summed across all images for each individual subject and then converted to micrometers and multiplied by the section thickness (40 μm) to calculate the volume. The estimates of the number of BrdU-positive cells were expressed as a density, the number of positive cells per cubic micrometer of the granular cell layer (BrdU-positive cells). For the Collagen IV microvasculature analysis, the granular cell layer and molecular layer Collagen IV staining density was analyzed in Fiji. First, the layers were outlined, then the images were subjected to the default auto-threshold in Fiji, and the area stained by Collagen IV was obtained. Estimates of the area stained by Collagen IV were obtained by dividing the area stained by the total area of the dentate gyrus.

### 2.5 Immunofluorescence and Image Analysis: Cell Differentiation

Differentiation rates of new (BrdU) cells into neurons and astrocytes was determined by staining a separate set of sections from each animal with primary antibodies against BrdU (1:200; AbCam Cat# ab6326), a marker of mature neurons, neuronal nuclei (NeuN, 1:400; Novus Biologicals Cat# NBP1-92693), and the marker of astrocytes S100 calcium-binding protein β (S100β, 1:1000; Synaptic Systems Cat# 287006(SY)). The staining protocol was identical to the above protocol with the omission of the ABC and DAB steps listed above. The free-floating sections were treated with secondary antibodies conjugated to fluorescent markers (DyLight 405 Jackson ImmunoResearch Labs Cat# 103-475-155; Alexa Fluor 647 Jackson ImmunoResearch Labs Cat# 112-605-003; Alexa Fluor 488 Jackson ImmunoResearch Labs Cat# 115-545-166) all at a 1:250 dilution. Images were captured with a Zeiss LSM 700 confocal microscope under a 40X oil objective. A total of 537 BrdU cells in the granule cell layer were analyzed for co-labeling with NeuN or S100β. Cells were sampled from all animals from both cohorts following standard procedures^59–62^.

### 2.6 Statistics

All statistical analyses were conducted using R (version 4.4.2; R Core Team). Data are presented as mean ± standard error of the mean (SEM). One-way analysis of variance (ANOVA) followed by Tukey’s post hoc test was used to assess differences in BrdU cell numbers and vascular area fraction measurements across the three treatment groups. Statistical significance was defined as *p* < 0.05. Normality and homogeneity of variance were assessed prior to performing parametric tests. Cell differentiation rates were compared between treatment groups using the chi-square test for independence.

## 3. Results

### 3.1 EV donors showed expected increases in cell genesis

We first quantified the total running distance of EV donor mice in two independent cohorts and found no significant difference between them (**Fig. 1B**). Mice in Cohort 1 ran an average of 323.9 km over 4 weeks, while mice in Cohort 2 ran an average of 394.5 km. To confirm that the exercised donor mice indeed displayed increased neurogenesis, we examined brains from a subset of Cohort 1 mice. BrdU immunostaining revealed a significant increase in BrdU-positive cells within the granule cell layer (GCL) of the hippocampus in runner mice (n = 4) compared to sedentary controls (n = 5; unpaired two-tailed *t*-test, *t*(7) = 12.44, *p* < 0.001) (**Fig. 1C**).

**Figure 1.**
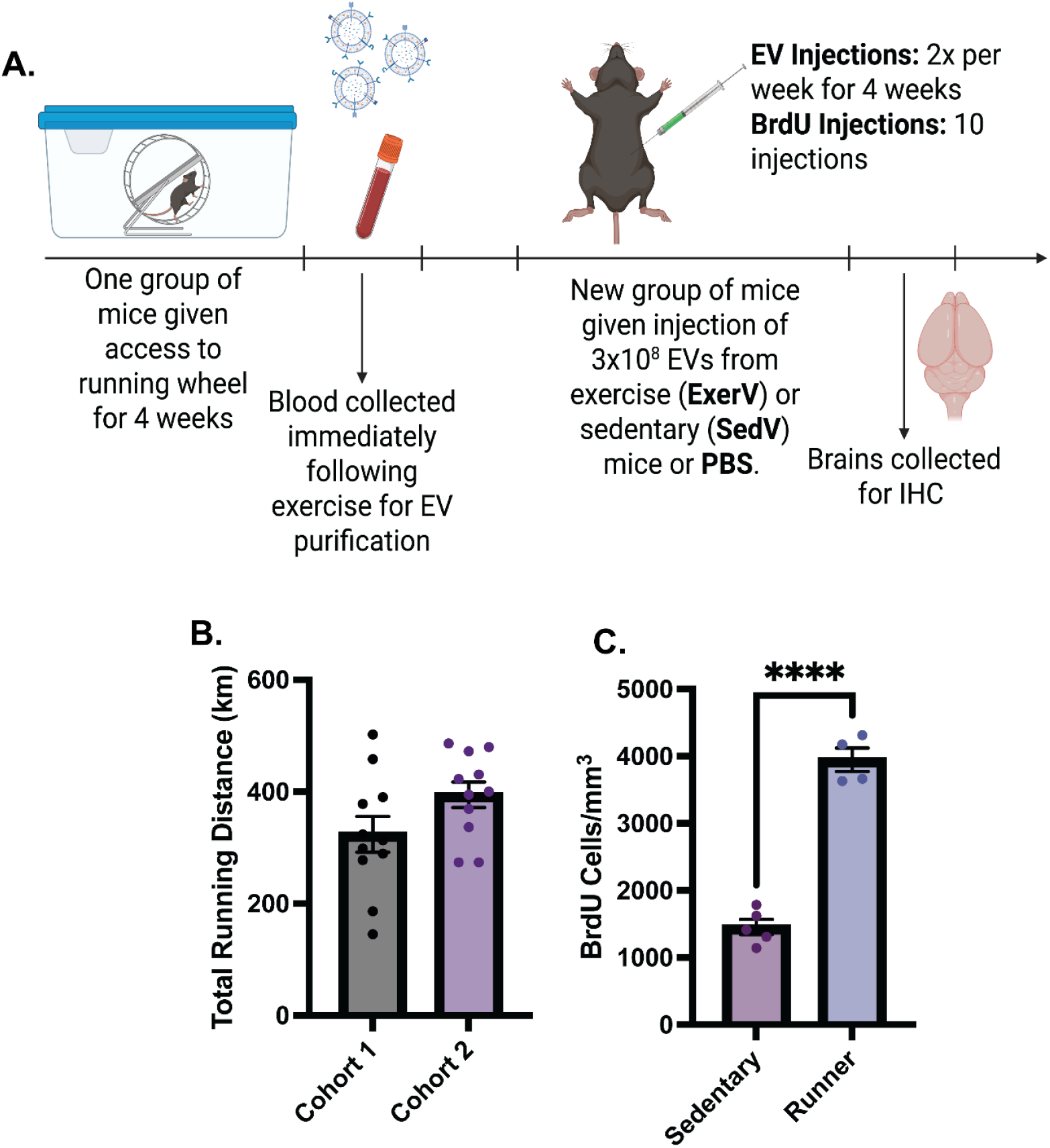
Experimental timeline and quantification of running behavior and BrdU+ cells. (A) Schematic timeline of the experimental paradigm. (B) Total running distance (km) over 4 weeks in two independent cohorts of donor mice (n = 10–11 per group). (C) Quantification of BrdU^+^ cells in the dentate gyrus of sedentary versus runner mice (n = 5 per group). Data shown as mean ± standard error of the mean (SEM); ****p < 0.0001, unpaired t-test.

### 3.2 Treatment with ExerVs enhanced cell genesis, neurogenesis, and astrogliogenesis

We assessed if administration of ExerVs could increase neurogenesis and astrogliogenesis in the GCL of the hippocampus. We found that ExerVs significantly increased the number of BrdU-positive cells in the GCL compared to both PBS- and SedV-treated mice (F(2, 26) = 3.639, p < 0.05) (**Fig. 2A**). Importantly, the effect was observed in an initial cohort, and was replicated in a second cohort, indicating that ExerVs increased cell genesis in the hippocampus. However, this was not accompanied by any changes in the volume of the GCL (p > 0.05) (**Fig. 2B**). To determine the proportion of new cells in GCL that differentiated into neurons or astrocytes, we sampled a total of 537 BrdU cells from all the animals and determined the percentage of BrdU cells co-expressing NeuN for new neurons or S100β for new astrocytes. Data showed no significant differences between groups in the differentiation rates (chi-square test, p=0.20). Collapsed across all three groups, approximately 90% of BrdU-positive cells were new neurons, 6% of BrdU-positive cells were new astrocytes, and 5% were new cells of undetermined phenotypes (**Table 1**).

**Table 1.**
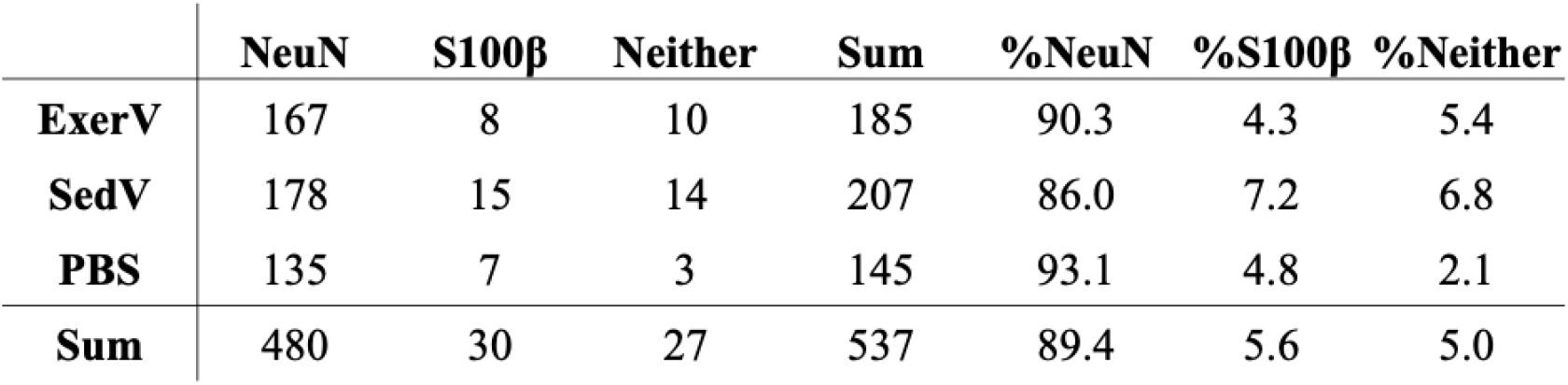
BrdU cell differentiation rates into neurons and astrocytes.

| | NeuN | S100β | Neither | Sum | %NeuN | %S100β | %Neither |
| --- | --- | --- | --- | --- | --- | --- | --- |
| **ExerV** | 167 | 8 | 10 | 185 | 90.3 | 4.3 | 5.4 |
| **SedV** | 178 | 15 | 14 | 207 | 86.0 | 7.2 | 6.8 |
| **PBS** | 135 | 7 | 3 | 145 | 93.1 | 4.8 | 2.1 |
| **Sum** | 480 | 30 | 27 | 537 | 89.4 | 5.6 | 5.0 |

**Figure 2.**
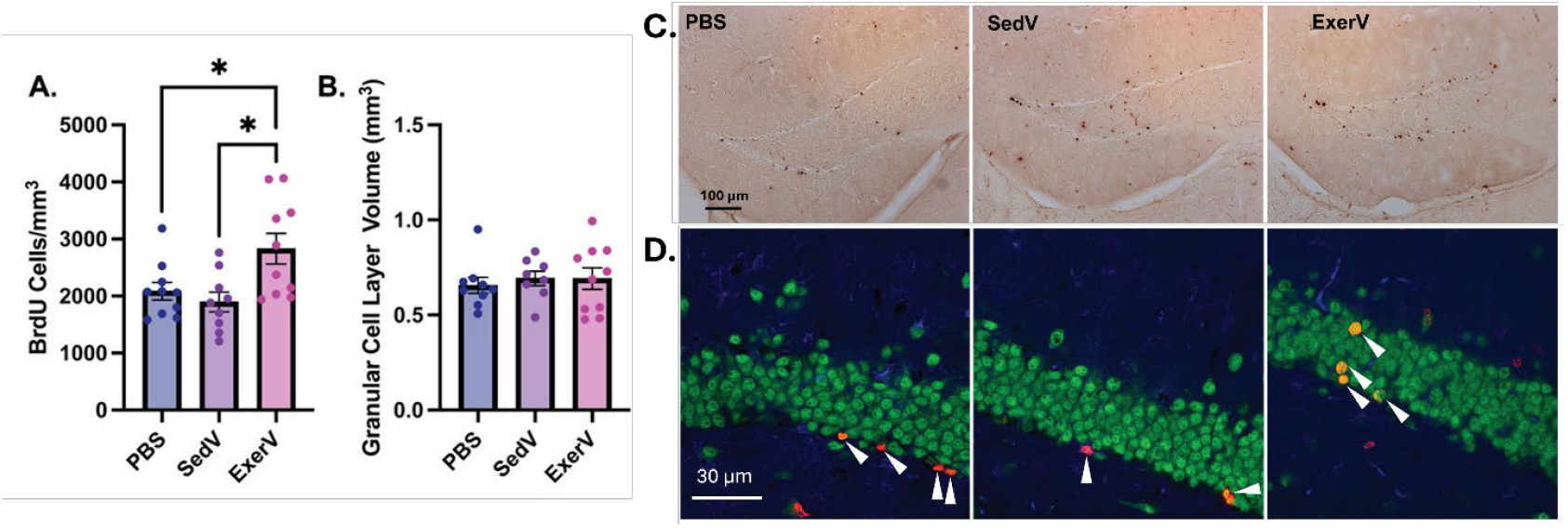
ExerV treatment increased neurogenesis and astrogliogenesis in the granular cell layer of the hippocampus. (**A & B**) Increased cell genesis was found in the GCL but this was not accompanied by an increase in GCL volume. * indicates a significant difference. Bars represent means ± SEM. (**C**) Representative images of BrdU-DAB staining for cell genesis are shown across the three treatment groups. (**D**) Representative images of the triple label fluorescent images taken to determine cell differentiation rates are shown. White flags pointing to new neurons (red/orange), and one new astrocyte in middle panel (pink). Approximately 89% of the BrdU cells differentiated into neurons and 6% into astrocytes with no differences between treatment groups (see Table 1).

### 3.3 ExerV treatment did not alter hippocampal microvasculature

To determine whether ExerV treatment increased supportive cellular components alongside neurogenesis in the hippocampus, we assessed vasculature using Collagen IV immunostaining. We hypothesized that ExerVs might promote increased vascular area via angiogenesis or hypertrophy of micro-vessels. However, quantification of vascular area revealed no significant differences across treatment groups in the GCL (**Fig. 3A**) or molecular layer (**Fig. 3B**) (p > 0.05).

**Figure 3.**
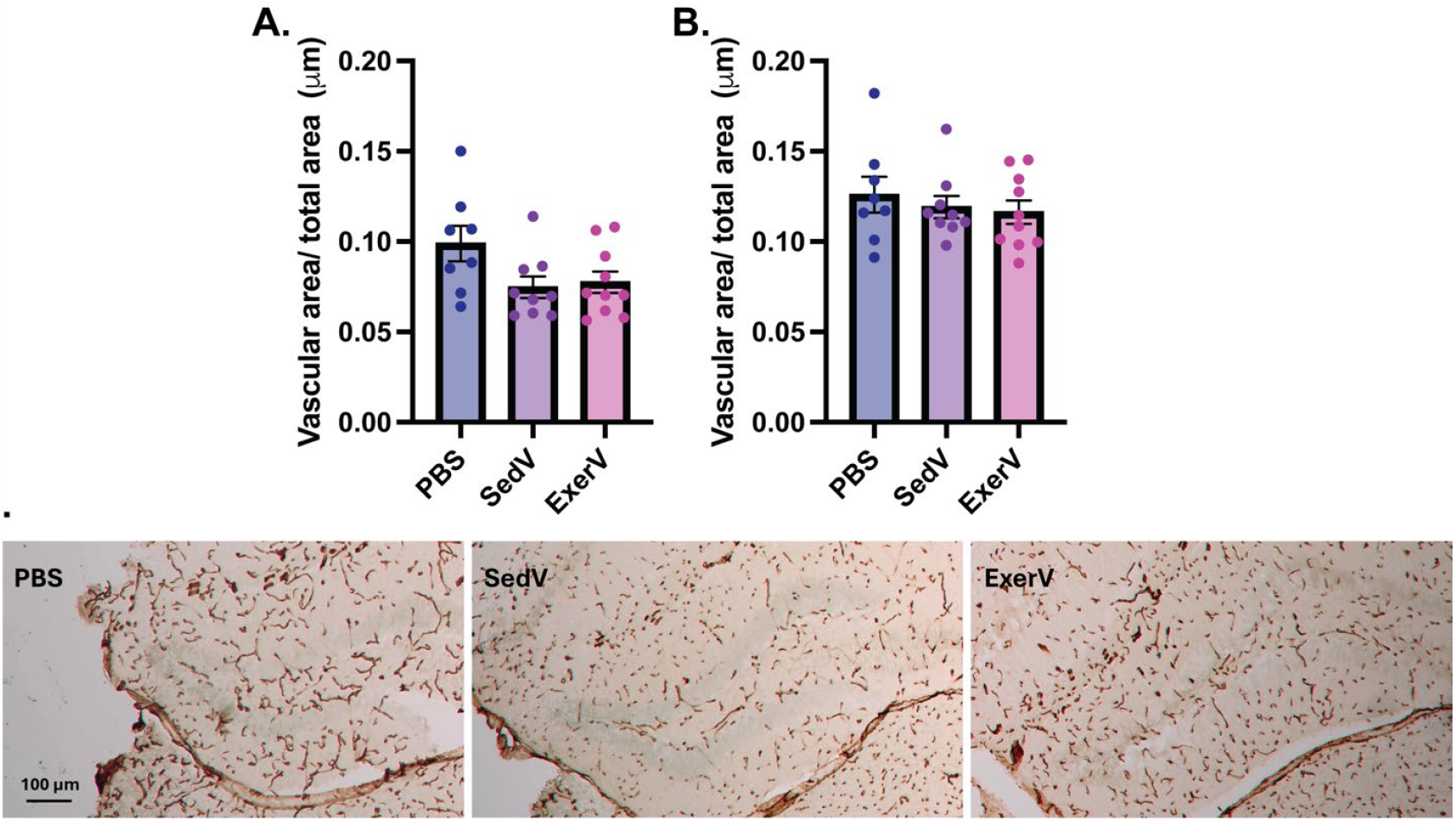
ExerV treatment did not alter vascularization in the hippocampus. (**A & B**) Quantification of vascular area relative to total area revealed no significant differences across treatment groups in the GCL (**A**) or molecular layer (**B**) in both cohorts. (**C**) Representative images of Collagen IV-DAB staining show vascular structures in hippocampal sections.

## 4. Discussion

Results suggest that extracellular vesicles (EVs) derived from exercising mice (ExerVs) significantly increase neurogenesis and astrogliogenesis in the dentate gyrus of sedentary mice. The result was replicated in two separate cohorts of mice, bolstering the rigor of the finding. No changes in hippocampal microvasculature were observed, suggesting that ExerVs selectively enhance neural precursor proliferation and differentiation without broadly altering the local cellular support environment. These results highlight a novel mechanism by which exercise may exert neurogenic effects through circulating vesicles that deliver molecular cargo to the brain.

One possible mechanism underlying the pro-neurogenic effect of ExerVs is the molecular composition of ExerVs. Proteomic profiling of ExerVs conducted by our collaborators^47^ revealed enrichment of proteins associated with neuroplasticity, cellular signaling, and synaptic remodeling. Among them, Spp1 (osteopontin) stood out as a multifunctional secreted glycoprotein known to modulate immune signaling pathways, enhance neuronal survival under stress conditions, and support the proliferation and integration of new neurons in the adult hippocampus ^48,49^. Also elevated was Msn (moesin), a member of the ERM family that links the actin cytoskeleton to membrane proteins—functions that are essential during neurite outgrowth, synapse stabilization, and the structural remodeling that underlies learning and memory ^50,51^. In addition, they observed increased expression of Psma3 and Psmb2, core subunits of the 20S proteasome complex, which mediate the selective degradation of damaged or short-lived proteins. Their role in maintaining synaptic proteostasis is particularly relevant in the context of exercise, which is known to enhance both protein turnover and neurogenesis in the hippocampus ^52,53^. The presence of these proteins offers a plausible mechanistic basis for the observed increases in neural proliferation and differentiation following ExerV treatment.

Contrary to expectations, however, ExerV treatment did not alter vascular density in key regions of the hippocampus. This may suggest that ExerVs enhance neurogenesis and astrogliogenesis through mechanisms that do not require concurrent vascular remodeling. Indeed, several neurotrophic factors elevated by exercise, such as brain-derived neurotrophic factor (BDNF) and insulin-like growth factor 1 (IGF-1), are known to promote neurogenesis independently of angiogenesis—raising the possibility that ExerVs similarly carry cargo capable of directly acting on neural precursor cells^6,13,18,63–66^.

Importantly, neither ExerV-treated mice nor the exercise donor mice themselves showed significant changes in dentate gyrus volume, despite robust increases in neurogenesis. Although physical exercise has been reported to increase DG volume, particularly in association with enhanced neurogenesis, these volumetric changes tend to be modest and are not consistently detectable across studies or methodologies^8,9^. Our findings are therefore not inconsistent with the existing literature and suggest that structural volume changes may lag behind or require a higher threshold of cumulative cellular and synaptic alterations to become apparent. Alternatively, ExerV-induced neurogenesis and even voluntary running-induced neurogenesis may be balanced by homeostatic mechanisms that maintain gross anatomical volume, such as synaptic pruning or glial remodeling, underscoring the complexity of interpreting volumetric outcomes in relation to cellular plasticity^67,68^.

Taken together, our data add to a growing body of evidence highlighting peripheral signals—particularly extracellular vesicles—as key mediators of brain plasticity^13,19,34–36^. The ability of ExerVs to promote neurogenesis and gliogenesis in sedentary mice supports the idea that exercise can confer its central nervous system benefits through transferable systemic factors. This raises promising therapeutic possibilities for leveraging EV-based approaches to mimic the cognitive and neuroprotective benefits of exercise, particularly for individuals who are unable to engage in regular physical activity due to age, injury, or illness. Given the central role of hippocampal atrophy and impaired neurogenesis in disorders such as depression, PTSD, and Alzheimer’s disease^69–71^, targeting the EV signaling axis may offer a novel, non-invasive strategy to restore or enhance brain plasticity in these vulnerable populations.

While this study provides compelling evidence for the neurogenic potential of ExerVs, several limitations warrant consideration. The current analysis was limited to male mice, and future studies will be needed to determine whether similar effects occur in females. Additionally, although we identified structural changes in neurogenesis and astrogliogenesis, behavioral correlates such as learning, memory, or affective outcomes were not assessed and remain to be explored. Finally, the primary cellular origin for EVs in plasma after exercise remains unknown. Despite these limitations, our results demonstrate that systemic delivery of exercise-derived EVs is sufficient to stimulate hippocampal plasticity, offering new insights into the molecular mechanisms of exercise and laying the groundwork for future translational applications.

## CRediT Author Contributions

**Meghan G. Connolly:** Conceptualization, Methodology, Formal Analysis, Software, Validation, Investigation, Writing – Original Draft, Writing – Review & Editing, Visualization, Project Administration. **Alexander M. Fliflet:** Conceptualization, Methodology, Formal Analysis, Software, Investigation, Writing – Original Draft, Writing – Review & Editing, Visualization, Project Administration. **Prithika Ravi:** Investigation. **Dan I. Rosu:** Investigation. **Marni D. Boppart:** Conceptualization, Methodology, Investigation, Writing – Review & Editing, Project Administration, Resources, Supervision, Funding acquisition. **Justin S. Rhodes:** Conceptualization, Methodology, Investigation, Writing – Review & Editing, Supervision, Project Administration, Resources.

